# Learned attraction replaces innate aversion in the evolution of an olfactory response in *Mus musculus*

**DOI:** 10.64898/2026.11.11.765886

**Authors:** Paul B. Conway, Zijun Wang, Nishan Shettigar, Esteban Urrieta, Íomar McCrea, Emily Dennis, Tomás J. Ryan

**Affiliations:** School of Biochemistry and Immunology, Trinity College Dublin, Dublin, Ireland; Trinity College Institute for Neuroscience, Trinity College Dublin, Dublin, Ireland; Janelia Research Campus, Howard Hughes Medical Institute, Ashburn, VA, USA; Florey Institute of Neuroscience and Mental Health, Melbourne Brain Centre, University of Melbourne, Melbourne, Victoria, Australia; Child & Brain Development Program, Canadian Institute for Advanced Research (CIFAR), Toronto, Ontario, Canada

## Abstract

The evolution of behavior is typically attributed to changes in innate circuitry, but learning can also drive behavioral evolution. *Mus musculus* evolved to produce trimethylamine (TMA) and behaviorally approach it, unlike the ancestral response retained by other mammals including *Mus caroli*. Here we demonstrate that this attraction is not innate but learned, depending on developmental TMA exposure. Whole brain immediate-early gene mapping reveals experience-dependent patterns of activation in response to TMA olfaction. Using activity-dependent TRAP2 labelling, we find that TMA-responsive neurons in the anterior olfactory nucleus are necessary and sufficient for attraction, consistent with a memory engram. Meanwhile, *Mus caroli’s* innate aversion depends on excitatory neurons in the posterior cortical amygdala. These results demonstrate how innate and learned circuitry both contribute to behavioral evolution in mammals.

## Introduction

Innate and learned behaviors are often treated as distinct, yet many behaviors depend on interactions between innate and learned circuits (*1–3*). How such interactions shape the evolution of behavior and neural circuit organization remains unclear (*4*, *5*). By taking a comparative approach across closely related species, it is possible to infer what changes have occurred in the brain over evolution to enable changes in behavior (*6*, *7*). Here we use the olfactory response to trimethylamine (TMA) in *Mus musculus* and *Mus caroli* as a model to investigate this question. TMA is a volatile amine associated with putrefaction and is broadly an aversive olfactory cue across mammals, including rats and humans (*8–10*). However, *M. musculus* have evolved to produce TMA in the urine of mature males and display an evolutionarily derived approach behavior to the odor. This relatively novel instinct has been demonstrated to play a role in social signaling (*9*, *11*, *12*). *M. caroli* is the most closely related species which does not produce TMA and is believed to retain the ancestral phenotypic aversive response (*9*, *10*).

Whether this evolutionary shift in valence reflects changes in innate circuits, recruitment of learning-dependent pathways, or broader circuit reorganization is unknown. Here, we compare behavioral responses and neural substrates of TMA olfaction across these species. We find that TMA attraction in *M. musculus* is experience-dependent and, by whole brain analysis of immediate-early gene expression, that whole brain responses, including recruitment of the hindbrain, thalamus and pallidum, are also experience-dependent. TRAP2 labelling reveals that TMA attraction is driven by the anterior olfactory nucleus. Meanwhile *M. caroli* aversion lacks these whole brain responses observed in *M. musculus*, and this aversion is instead driven by the cortical amygdala. These results illustrate how evolutionary changes can involve changes to both innate and learned responses to odor which function in a complementary manner to enable adaptive behavior.

## Results

### TMA attraction in *musculus* is learned and sex-specific, while *caroli* innately avoids TMA

To test for the presence of TMA, we performed ^1^H-NMR analysis on urine from male and female *M. musculus* and *M. caroli.* In line with previous reports (*9*), we demonstrated that male *M. musculus* produce TMA in their urine, while it was not detectable in female *M. musculus* or either sex of *M. caroli* (Fig. 1 A-B). Dietary administration of the antibiotic doxycycline to deplete the microbiome results in significant reduction of TMA levels from male mouse urine (Fig. 1 B-C and Fig. S1 J-K). To assess behavioral responses of these two species to TMA, an olfactory arena was constructed based on designs from Root et al. (*13*), to control odor delivery in both a spatial and temporal manner (Fig. 1 D and Fig. S1 A-G). Odor was delivered with concentration increasing approximately 30 seconds after switching of valve status and plateauing after approximately 3 minutes as measured by a photoionization detector (Fig. S1 A-B). By delivery of odor to specific ports, it is possible to induce behavioral attraction (Fig. 1 E-G) or aversion using the predator odor 2,4,5-trimethylthiazoline (TMT) (Fig. S1 D-G) to specific quadrants. To determine which concentration of TMA is most attractive to *M. musculus*, a concentration assay was performed, and peak attraction to TMA was observed at 184 mM (Fig. S1 C). *M. caroli* were then tested at this same concentration and demonstrated an aversive response to TMA (Fig. 1 H-L). When tested in a simple arena with droplets of TMA placed at either end, the same species-specific responses were observed (Fig. S1 H-I). These results demonstrated that *M. caroli* retain the ancestral behavioral phenotype of TMA aversion.

**Fig 1.**
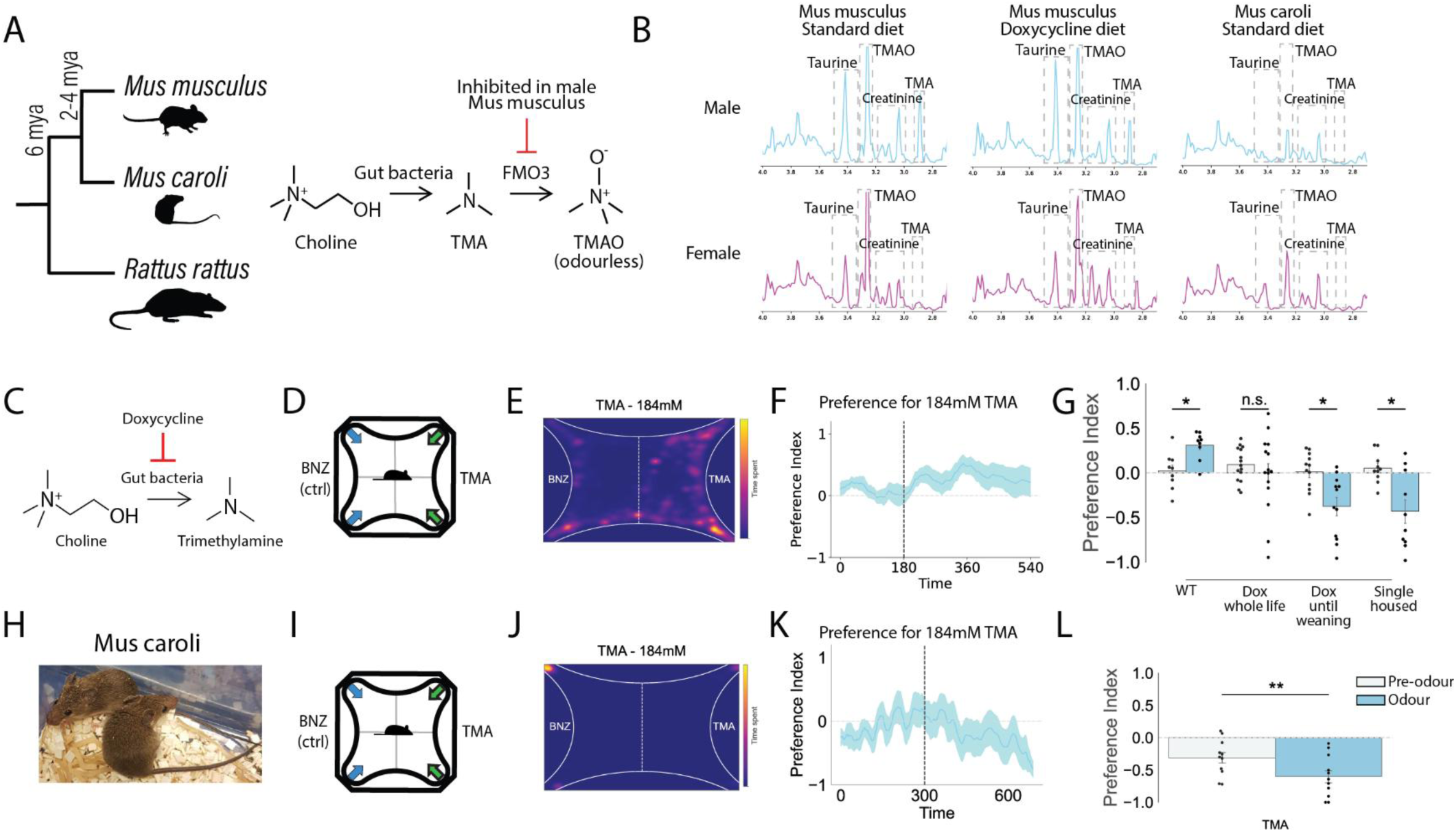
*Mus musculus* demonstrate experience-dependent attraction to trimethylamine. (A) Phylogeny of *Mus musculus, Mus caroli* and *Rattus rattus*. Gut bacteria produce TMA in all species, but in male *musculus* the FMO3 enzyme which oxidizes TMA is inhibited. (B) ^1^H-NMR spectra for urine from male and female *Mus musculus* and *Mus caroli*, and male and female *Mus musculus* on doxycycline diet. (C) Doxycycline administration kills gut bacteria, preventing production of trimethylamine. (D) Schematic of vacuum-powered odor delivery apparatus. (E) Heatmap of TMA response (n = 9) at 184 mM. (F) Timetrace indicating how odor preference index evolves over time (n = 9, shaded area represents standard error of the mean). Odor presentation indicated by a vertical dashed line. (G) Response of *Mus musculus* to 184 mM TMA under normal conditions (n = 9), whole-life doxycycline diet (n = 15), doxycycline diet until weaning (n = 12) and chronic single housing (n = 10). (H) Image of 2 *Mus caroli* males. (I-K) As in D-F, but for *Mus caroli* (n = 11). (L) *Mus caroli* preference index for 184 mM TMA (n = 11). n.s. *p* ≥ 0.05, \**p* < 0.05

As *M. musculus* produce TMA, their response to TMA could be learned, innate, or both. To investigate which, TMA production in *M. musculus* was manipulated and the behavioral response of mice raised in a TMA-depleted environment was tested. TMA is produced by commensal microflora in the mouse gut, and antibiotic administration reduces the amount of TMA present in mouse urine (*9*). A diet containing doxycycline was administered to the mice, resulting in a decrease in TMA concentrations in male mouse urine (Fig. 1 B and Fig. S1 K) and a loss of TMA attraction, even when only administered until weaning (Fig. 1 G). This suggests TMA attraction is learned and depends on TMA exposure during the juvenile period. To investigate the social signaling role of TMA, mice were chronically single housed from weaning, a procedure known to induce social aversion (*14*, *15*), and TMA responses were assessed. Socially isolated mice demonstrated aversion to TMA, indicating that TMA responses may be influenced by social drive (Fig. 1 G). Additionally, as the production of TMA in *M. musculus* is sex-specific, the response of female mice to TMA was assessed. On average, female mice show no preference to TMA (Fig. S1 L). However, when the responses of female mice are separated on the basis of their estrus cycle, female mice show TMA avoidance during metestrus/diestrus, while being neutral or mildly attracted to TMA when in estrus/proestrus (Fig. S1 M). This supports a sex-specific social signaling role of TMA. To determine which brain areas may be involved in the TMA olfactory response, c-Fos expression patterns in the olfactory cortex were investigated following TMA exposure, identifying an increase of c-Fos expression in the anterior olfactory nucleus (AON) in *M. musculus*, while the posterior cortical amygdala showed a decrease (Fig. S1 N).

### Whole brain light sheet imaging reveals global changes in neural responses to TMA between *Mus musculus* and *Mus caroli*

The above results demonstrate that *M. caroli* innately avoid TMA while *M. musculus* demonstrate experience-dependent TMA attraction. However, as the odor may have other associations beyond pure olfactory perception in both species, such as social signaling in *M. musculus* or disgust in *M. caroli*, it is likely that more brain areas are involved. To explore this possibility, whole brain immediate-early gene expression following odor exposure was investigated. Mice were exposed to either TMA or H2O in the olfactory arena for 5 min and perfused 60 min later. Brains were then cleared and stained for both c-Fos and Arc using an iDisco protocol (*16*) and imaged via light sheet microscopy (Fig. 2 A). Raw images were then transformed into a common coordinate framework (CCF) based on the Allen Atlas (Fig. 2 B) using code developed by Dennis et al. (*17*), with automated cell counting performed via ClearMap following manual parameter calibration to match cell detection rates with that of two human annotators (*18*). This was performed for male *M. musculus*, dox-fed male *M. musculus*, female *M. musculus* and male *M. caroli*, exposed to either clean air or TMA. Broadly, TMA olfaction resulted in an increase in c-Fos expression in the pallidum, striatum, thalamus, hypothalamus and hindbrain in standard diet male *M. musculus* (Fig. 2 C-D), but not dox-diet males (Fig. 2 G-H).

**Fig 2.**
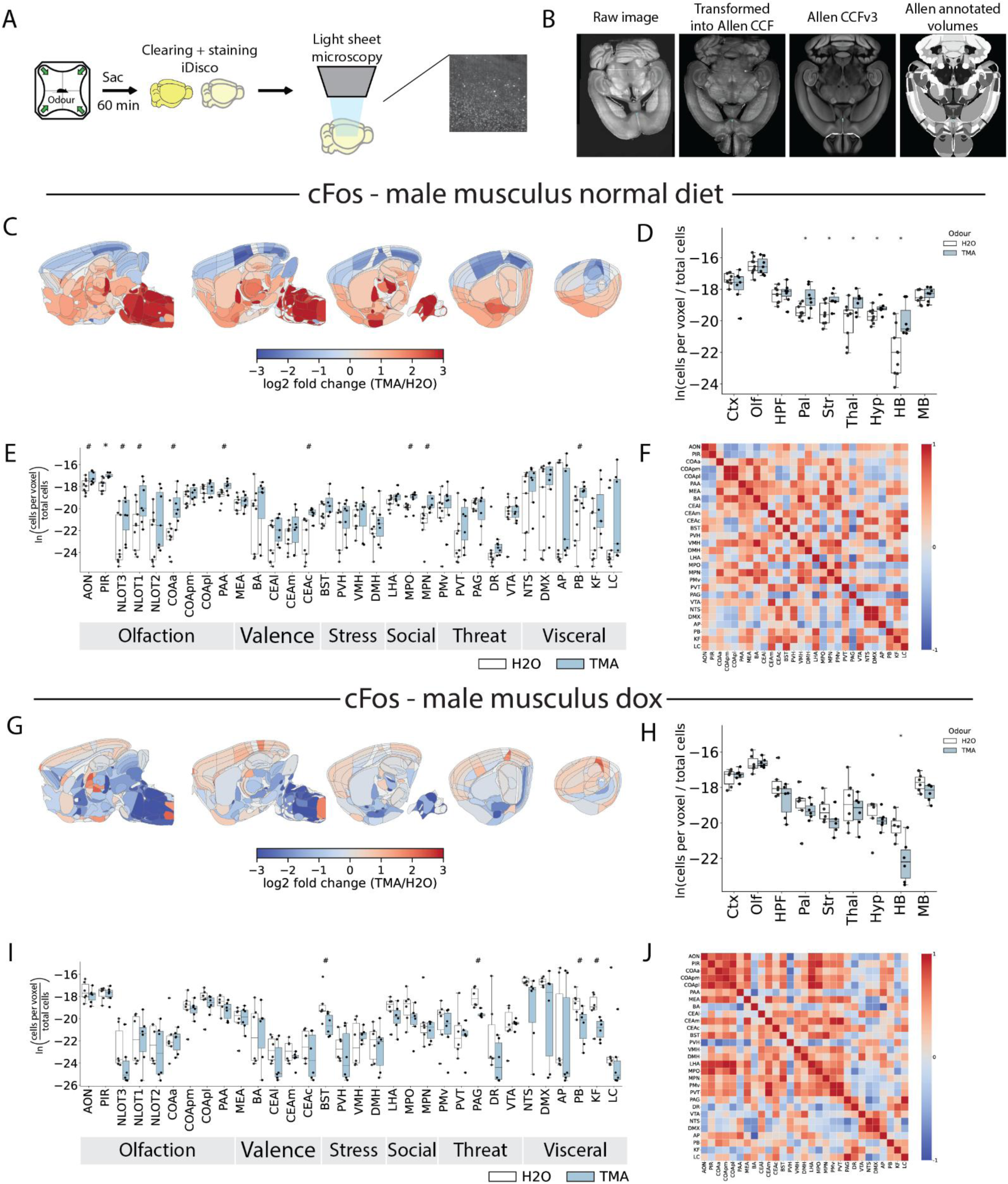
Increased c-Fos expression in the olfactory cortex and hindbrain in standard diet but not doxycycline diet *Mus musculus.* (A) Brains were cleared and stained according to the iDisco protocol and imaged on a light sheet microscope. (B) Raw images were transformed into a common coordinate framework and then fixed to the Allen Atlas template. Cells within each Allen Atlas region are quantified. (C) Fold change in c-Fos expression in the whole brain during TMA exposure (n = 7) vs. H2O exposure (n = 9) for male *Mus musculus* on a standard diet. (D) Cell counts for TMA and H2O exposed mice. (E) Relative c-Fos expression levels in selected brain regions during TMA vs. H2O exposure for male musculus mice on a standard diet. (F) Correlations of c-Fos expression across selected brain regions for TMA-exposed male musculus mice on a standard diet. (G-J) As in C-F, but for male *Mus musculus* on a doxycycline diet (H2O n = 6, TMA n = 6). *# p (*uncorrected) < 0.05, * q (FDR corrected p value) < 0.05

We hypothesized that *M. musculus* have lost putrefaction-linked TMA aversion over the course of evolution, due to altered activity in brain areas associated with attraction and social behavior. To test this prediction, brain areas associated with these behaviors were selected for closer inspection based on known involvement in: olfaction (*19*, *20*), odor valence (*13*, *21*), stress (*22–24*), social behavior (*25–27*), threat (*28–30*), and visceral/interoceptive signaling (*31–34*).

Male *M. musculus* show increases in c-Fos expression in many of the selected areas, including AON, PIR, NLOT, PAA (olfactory areas), CEAc (implicated in valence and stress regulation), MPO, MPN (social behaviors) and PB (visceral and interoceptive signaling) (Fig. 2 E-F). Furthermore, there were positive correlations in c-Fos expression between these areas in male *M. musculus* exposed to TMA, with a negative correlation between the AON and the cortical amygdala (Fig. 2 F and Fig. S6). All of these TMA-induced increases were absent in female *M. musculus* (Fig. S2 E-F) and dox-fed male *M. musculus* (Fig. 2 I-J), which are both neutral in their response to TMA. Meanwhile, there was a positive correlation between c-Fos expression in the AON with the cortical amygdala in dox-fed *M. musculus* (Fig. 2 J). In females, decreases were observed in MEA (social and sex-related chemosensory signals), CEAm (aversion), PVT (anxiety/stress), PAG and DR (threat), with dox-fed males showing decreases in BST (social stress), PAG (threat), PB and KF (visceral and interoceptive signaling). For Arc, there are overall fewer changes in activity across the three groups. In *M. musculus*, an increase in Arc expression is observed in the NTS (visceral and interoceptive signaling), with no changes in Arc expression levels observed in female *M. musculus* and dox-fed males (Fig. S3 A-L and Fig. S4).

Overall, these results indicate an increase in activity in *M. musculus* in areas related to olfactory attraction and social behavior, which is absent in mice raised on a doxycycline diet and in females who do not demonstrate TMA attraction. As such, these changes may be driven by a TMA-associated memory engram which drives TMA attraction.

### TMA-responsive AON cells are necessary and sufficient for attraction in male *Mus musculus*

Whole brain c-Fos quantification revealed an increase in c-Fos levels and correlation in various parts of the olfactory cortex, including the anterior olfactory nucleus (AON), while having a negative correlation with the cortical amygdala. The AON has been demonstrated to store olfactory memory engrams (*20*), including social memory (*35*). Therefore the AON, in addition to the posterior cortical amygdala (CoAp) which has been implicated in olfactory valence (*13*, *21*), were selected for further investigation.

To determine whether TMA-responsive cells in these two regions are necessary for TMA attraction, TRAP2 mice were injected with an AAV-DIO-hM4D(Gi)-mCitrine vector in either the AON or CoAp, and 4-hydroxytamoxifen (4-OHT; 50 mg/kg) administered following odor exposure to capture cells active during TMA olfaction (Fig. 3 A-B). For the AON, mice administered with the DREADD agonist clozapine-N-oxide (CNO) showed a slightly aversive response to TMA, while control mice administered saline remain attracted to TMA (Fig. 3 C-D). For the CoAp, mice remain attracted to TMA with saline administration, but become neutral towards TMA with CNO administration (Fig. 3 E-F). Therefore TMA-responsive cells in both the AON and CoAp are necessary for behavioral attraction to TMA. Meanwhile, for mice injected into the main olfactory bulb, inhibition of TMA-responsive cells induces behavioral aversion to TMA, mimicking what was observed for the AON (Fig. S7).

**Fig 3.**
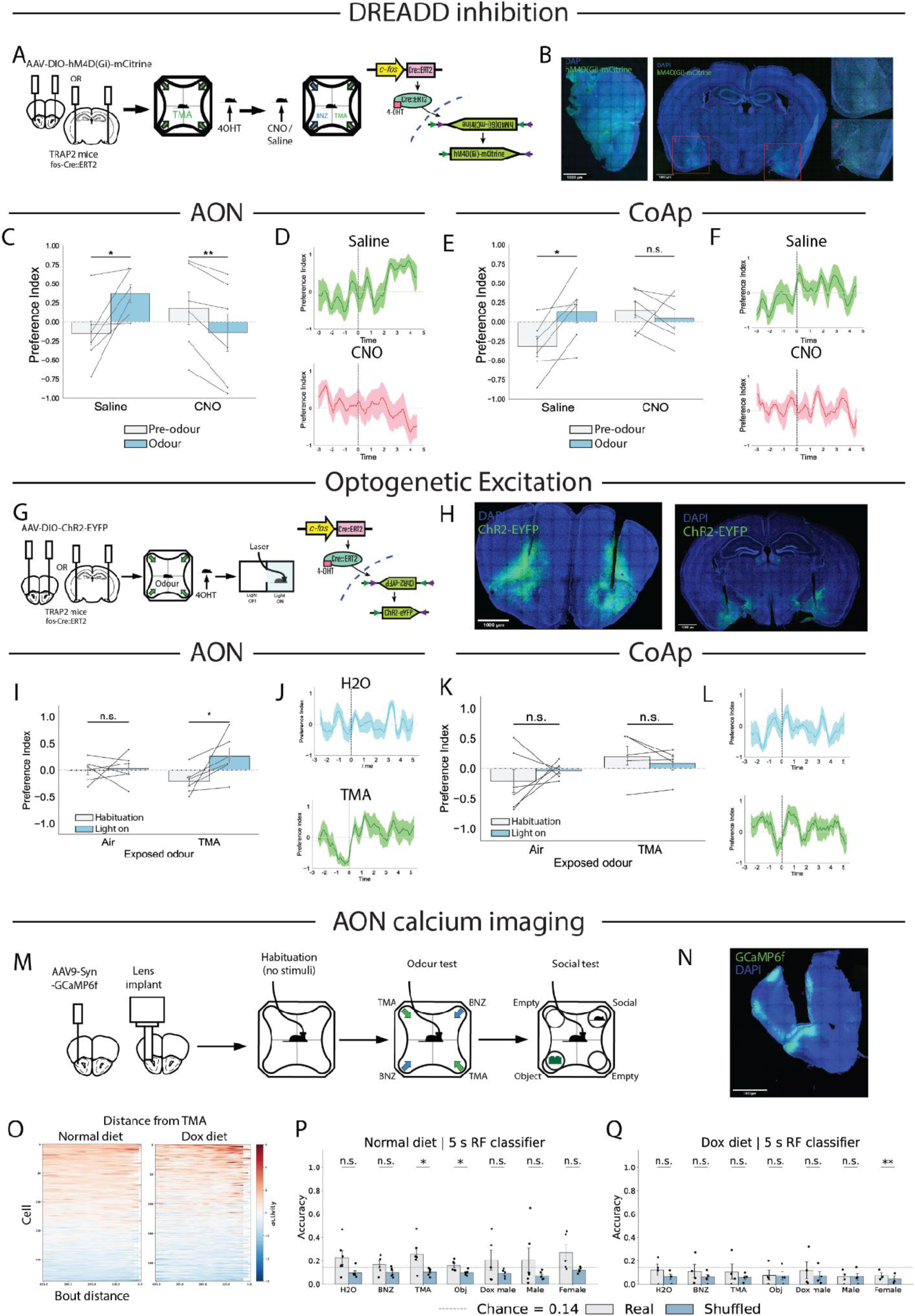
TMA-responsive ensembles in the AON drive attraction in *musculus,* while CoAp ensembles are necessary but not sufficient. (A) Schematic indicating behavioral timeline for engram labelling and engram inhibition during TMA olfaction. (B) Example AON slice showing viral expression. (C) Effect of no inhibition (saline) versus inhibition (CNO) of AON cells tagged during exposure to TMA (Student’s paired t test; saline, n = 6; CNO, n = 6). (D) Timetraces indicating average preference index across mice for saline (top) or CNO (bottom) administration. Odor onset indicated by dashed line. (E-F) As in C-D, but for the CoAp (n = 6). (G) Schematic indicating behavioral timeline for engram labelling and spatially-controlled engram reactivation. Laser stimulation occurs in one half of the arena only. (H) Example slices showing viral expression and implant track in the AON and CoAp. (I) Preference index pre- and post-onset of laser stimulation for AON cells tagged during exposure to clean air or TMA (Student’s paired t test; Air, n = 6; TMA, n = 6). (J) Timetrace indicating average preference index across mice during optogenetic stimulation of AON cells tagged during delivery of clean air (top) or TMA (bottom) odorized air. Laser onset phase indicated by dashed line. (K-L) As in I-J, but for the CoAp (Air n = 6, TMA n = 5). (M) Schematic indicating calcium imaging timeline. Implanted mice were recorded in the presence of no stimuli, odor stimuli or social stimuli. (N) Example slice indicating viral expression and implantation in the AON. (O) Heatmap indicating the average response of each cell (approximately 250 cells across 4 - 6 mice in each group) during approach to the stimulus location. (P) Performance of a random forest classifier on real vs. shuffled data from standard diet mice for each stimulus (n = 6). (Q) Performance of a random forest classifier on real vs. shuffled data from doxycycline diet mice for each stimulus (n = 4). n.s. *p* ≥ 0.05, \**p* < 0.05, \*\**p* < 0.01

To test the sufficiency of TMA-responsive cells, TRAP2 mice were injected with AAV-DIO-ChR2-EYFP in either the AON or the CoAp, and following odor exposure (clean air or TMA) 4-OHT was administered to allow expression of ChR2-EYFP in TMA-responsive cells (Fig. 3 G-H). Mice were then placed into a simple rectangular arena, in which optogenetic stimulation was delivered while the mouse was in one half of the arena only. For the AON, stimulation of cells captured during TMA olfaction, but not clean air, was sufficient to induce attraction to the light-ON component of the arena (Fig. 3 I-J). For the CoAp, stimulation of neither H2O nor TMA-TRAPed cells induced attraction to the light on component (Fig. 3 K-L). To determine whether these responses are general to the AON and CoAp, the predator odor 2,4,5-trimethylthiazoline (TMT) was also used (Fig. S8). Optogenetic stimulation of TMT-tagged cells in the AON resulted in no change in place preference, while stimulation of TMT-tagged cells in the CoAp resulted in avoidance of the light on portion of the arena (Fig. S8 A-D).

These results show that the TMA-responsive cells in the AON are both necessary and sufficient for learned TMA attraction in *M. musculus*, while CoAp TMA-responsive cells are necessary but not sufficient to drive attraction. This indicates a role for TMA-responsive ensembles in the AON in processing TMA-related odor identity, with the CoAp TMA-responsive ensembles being required for non-TMA specific olfactory valence processing. Based on our findings in Figure 2, doxycycline-fed mice should not possess such a TMA olfactory engram. To test this hypothesis, *in vivo* calcium imaging of the AON was performed on mice fed either a standard diet or doxycycline diet for their whole lives (Fig. 3 M-N). While both groups of mice contained ensembles of cells which were reliably responsive to TMA (Fig. 3 O), only in standard diet mice was a random forest classifier capable of distinguishing true TMA data from shuffled data (Fig. 3 P-Q). This suggests that doxycycline mice lack the AON olfactory engram for TMA, which is required to drive TMA attraction.

### *Mus caroli* show distinct patterns of c-Fos expression in the whole brain with aversion driven by the CoAp

Thus far, these results demonstrate that *M. musculus* TMA attraction is driven by the AON, while *M. caroli* are repelled by TMA. To determine how this distinct behavior may have formed through evolution, whole-brain immediate early gene expression following odor exposure in *M. caroli* was investigated (Fig. 4 A-F). *M. caroli* lacked the broad increases in umbrella regions detected in *M. musculus* (Fig. 4 C-D), with the main olfactory area upregulated being the CoAp (Fig. 4 E). There were increases in c-Fos in the COApm (aversion area) and decreases in the PVT (anxiety/stress area) and PAG (threat area). Similarly to dox-fed *M. musculus*, the AON and the cortical amygdala showed a positive correlation in c-Fos expression in *M. caroli* exposed to TMA (Fig. 4 F and Fig. S5). For Arc, there was a general decrease in expression in the cortex, olfactory cortex and hypothalamus. There were specific increases in Arc expression in the nucleus of the olfactory tract and basolateral amygdala (Fig. S4 and Fig. S6).

**Fig 4.**
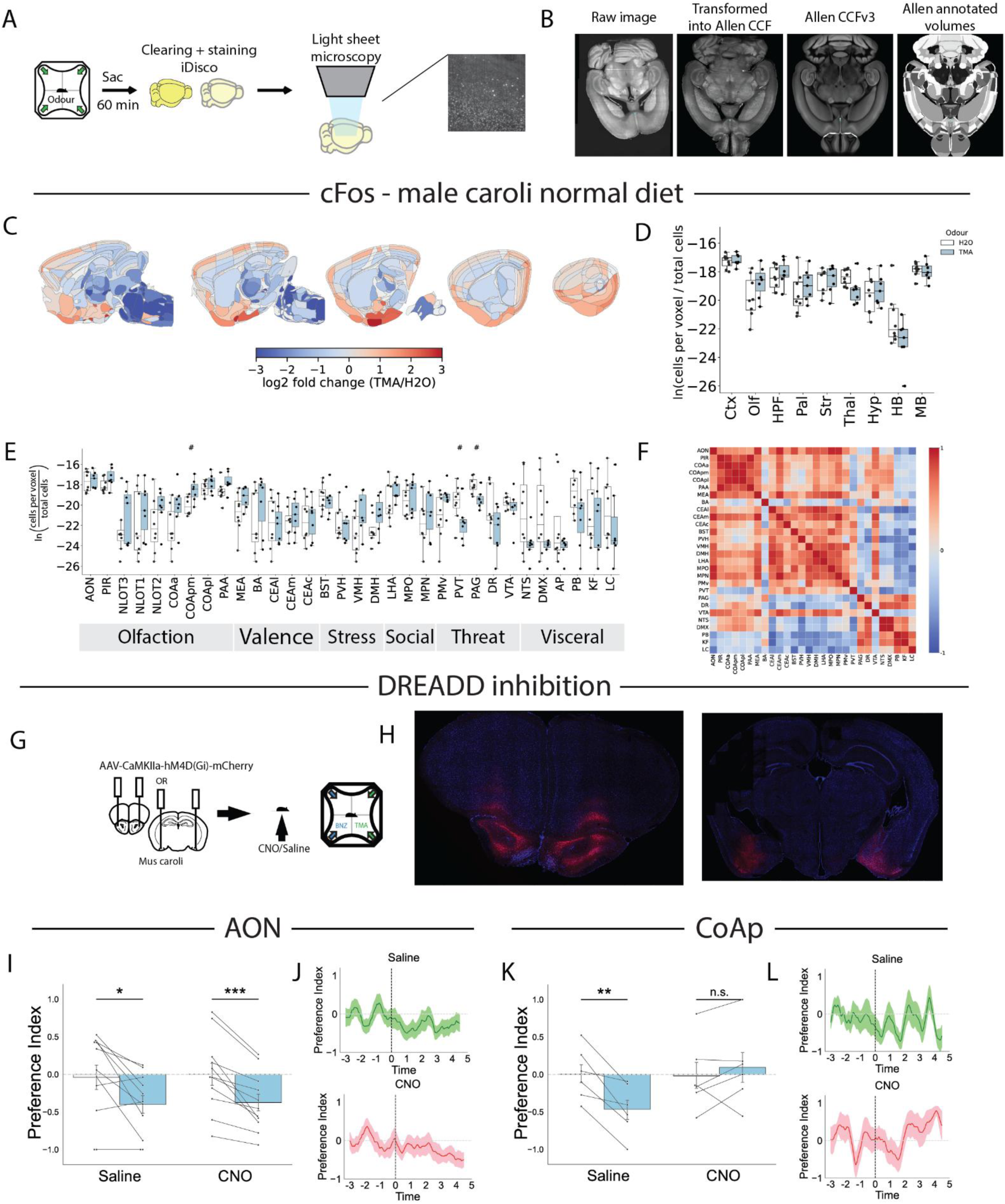
TMA aversion in *Mus caroli* is dependent on the CoAp and drives distinct c-Fos patterns across the whole brain. (A) Brains were cleared and stained according to the iDisco protocol and imaged on a light sheet microscope. (B) Raw images were transformed into a common coordinate framework and then fixed to the Allen Atlas template. Cells within each Allen Atlas region are quantified. (C) Fold change in c-Fos expression in the whole brain during TMA exposure (n = 7) vs. H2O exposure (n = 8) for male *Mus caroli*. (D) Cell counts for TMA and H2O exposed mice. (E) Relative c-Fos expression levels in selected brain regions during TMA vs. H2O exposure for male *caroli* mice (# *p* < 0.05, * q < 0.05). (F) Correlations of c-Fos expression across selected brain regions for TMA-exposed male *caroli* mice. (G) hM4D(Gi) was expressed under the control of the CAMKIIa promoter via viral infection in either the AON or CoAp. (H) Example AON (left) and CoAp (right) slices showing viral expression. (I) Effect of no inhibition (saline n = 11) versus inhibition (CNO n = 11) of AON CaMKIIa+ cells. (J) Timetrace indicating a 30s average preference index across mice with saline administration (top n = 11) and CNO administration (bottom n = 11). Odor onset indicated by dashed line. (K-L) As in I-J, but for the CoAp. n.s. *p* ≥ 0.05, \**p* < 0.05, \*\**p* < 0.01, \*\*\**p* < 0.001.

These results demonstrate a different broad response in the *M. caroli* whole brain compared with *M. musculus*. Specifically, an increase in the CoAp was detected without a change in activity in the AON. To determine whether these correlative evolutionary changes are functional, AAV-CaMKIIa-hM4D(Gi)-mCherry was injected into the AON or CoAp to express the inhibitory DREADD in CaMKII+ neurons (Fig. 4 G-H). For the AON, *M. caroli* retained TMA aversion regardless of CNO or saline administration (Fig. 4 I-J). For the CoAp, CNO administration resulted in a loss of TMA aversion, while control mice administered saline retained the aversive response (Fig. 4 K-L). These results show that CaMKII^+^ excitatory neurons in the CoAp but not the AON are required for TMA aversion in *M. caroli*. Overall, these results delineate an evolutionary shift with ancestral, innate CoAp-driven TMA aversion in *M. caroli* and derived, learned AON-driven and socially-related TMA attraction in male *M. musculus*.

## Discussion

We present an integrative study combining behavioral analysis, whole brain activity profiling, localized calcium imaging, engram labelling and manipulation, and cross-species analysis to investigate the evolutionary shift in TMA valence in the rodent lineage. *M. musculus* attraction to TMA arises through suppression or reweighting of ancestral aversion circuits alongside recruitment of learning to form a memory engram which drives socially-driven attraction. While *M. caroli* demonstrate innate CoAp-dependent aversion to TMA, attraction in *M. musculus* requires experience and depends on activity in the AON. The behavioral results also suggest that TMA functions as a social contextual signal in *M. musculus* rather than as a fixed attractive odor. Attraction is abolished by reducing developmental TMA exposure and altered by social isolation and reproductive state, indicating that behavioral valence depends on both sensory experience and internal state. This organization may allow flexible assignment of social meaning to a cue that is reliably encountered during development.

Whole-brain activity mapping suggests that the evolutionary transitions in TMA responsiveness involve distributed network-level changes, in addition to the specific changes which were functionally investigated. In *M. musculus*, TMA preferentially recruits regions implicated in olfactory association and social behavioral processing, such as the AON, PIR and MPO. In *M. caroli*, there is engagement of areas linked to innate olfactory aversion and avoidance, such as the CoApm and BA. The absence of many of these activity patterns in TMA-deprived *M. musculus* supports the interpretation that developmental experience reshapes large-scale neural responses to the odor. Together, these findings suggest that evolutionary changes in behavioral valence can emerge through reorganization of interactions between innate and learned circuits across the brain.

It has been proposed that circuits in the brain may be categorized as innately assembled, partially assembled but requiring some learning, or highly plastic (*1*). However, even innately assembled circuits may be subject to experience-dependent plasticity (*3*, *36*, *37*). Under this framework, these findings suggest an evolutionary switch in *M. musculus* from an ancestral state involving innately assembled circuits driving aversion, to a partially assembled circuit which relies on experience to link the neural circuit with relevant features of the developmental environment. Given the complexity of social behavior, which can require many different responses to the same stimulus depending on context, such a system would allow the brain to form a rich and complex representation of the sensory stimulus and allow adaptation of behavior in a context-appropriate manner based on the developmental environment. This evolutionary trajectory may represent iterated instances of the Baldwin effect (*38*), in which phenotypic plasticity and genetic assimilation alternate across successive evolutionary steps. An initial, developmentally acquired reduction in TMA aversion through TMA exposure may have become genetically assimilated to innate neutrality in *M. musculus*, producing a permissive baseline from which an AON memory engram for TMA attraction can form.

## Materials and Methods

### EXPERIMENTAL MODEL DETAILS

#### Animals

All *Mus musculus* behavior experiments and tissue harvesting were performed with C57BL/6J mice. TRAP2 mice were used for optogenetic and chemogenetic experiments (*39*, *40*). All *Mus caroli* experiments were performed using CAROLI/EiJ mice. Mice were housed in groups of 3-6 in cages with enrichment, food and water ad libitum under a 12 h light/dark cycle. The animal room was kept at a constant temperature of 22°C and 50% humidity. Mice used for experiments were 8-12 weeks of age. All experiments were performed during the light phase. All experimental manipulations were carried out in accordance with Health Products Regulatory Authority (HPRA) Ireland guidelines and the principles of the 3R’s (https://nc3rs.org.uk/) and with ethical approval from the Trinity College Dublin ethics committee and the Animal Research Ethics Committee from the HPRA.

| Strain | Jax ID |
|---|---|
| C57BL/6J | # 000664 |
| Fostm2.1(icre/ERT2)Luo/J (TRAP2) | # 030323 |
| CAROLI/EiJ | # 000926 |

## METHOD DETAILS

### 4-corner olfactory behavioral assay

The four-field behavior chamber was designed to replicate the four-field behavior chamber used by other groups (*13, 17*), and inspired by similar experiments performed in Drosophila (VET, LENTEREN et al. 1983, Faucher, Forstreuter et al. 2006, Semmelhack and Wang 2009). The chamber was an enclosed four quadrant arena with airflow out from the center powered by a vacuum (KNF Laboport® mini-pump). Airflow at 230 ml/min was pumped into each quadrant via gas-mass flow controllers (Omega). Airflow exited the chamber via a 20 mm outlet in the center of the floor. The chamber had a 76 cm diameter, a corner arc of 1.9 cm radius and a chamber height of 7.6 cm. The floor and the walls were made from polypropylene and the ceiling was clear, 1 cm thick acrylic. Odor was applied by solenoid valves redirecting airflow through 100 ml glass bottles containing 10 ml of diluted odorant, dissolved in deionized water. Behavior was recorded via a webcam mounted above the chamber. Custom software written in BONSAIrx, a reactive programming language (Lopes, Bonacchi et al. 2015), controlled valve position, tracked the position of the mouse in real time at a rate of 30 Hz, and synchronized this information for writing to csv. The behavior was further analyzed offline using custom scripts written in python. Mice were placed in the chamber for 8 min experiments and tested only once per day. The first 3 min served as a baseline test for bias within the arena. The following 5 min were used to assess behavioral response to the odor. To control for any spatial bias, animals were always tested with a 5 min period of no odor. Further, the area to which odor was delivered was altered across experiments. Preference index was calculated as the difference between the time spent in the test odor quadrants vs. control quadrants, divided by the total time of odor presentation.

### Doxycycline diet

To generate mice with reduced developmental TMA exposure ("whole-life" doxycycline cohort), standard chow was replaced with doxycycline-containing diet (40mg/Kg; LBS Serving Biotechnology Ltd) in breeding pairs, and pairs were maintained on doxycycline diet throughout mating, gestation and lactation. Offspring were maintained on doxycycline diet after weaning and used for experiments. For the “dox until weaning cohort”, offspring were provided with standard chow after weaning.

### Single housing

Mice in the single-housing cohort (Fig. 1G) were separated from littermates at weaning and housed individually until behavioral testing.

### Estrus cycle staging

Estrus cycle stage was determined by visual inspection of the vaginal opening, and classified as a binary readout (metestrus/diestrus vs. proestrus/estrus) rather than the full four-stage cycle.

### Simple olfactory behavioral assay (droplet test)

A box with a sealed lid had two squares of 1cm x 1cm filter paper placed at each end. 20µl of 184 mM TMA or 1 mM BNZ, dissolved in deionized water, was placed at each end of the arena. Animals were then placed into the arena, and their behavior was recorded via a camera directly above the arena. The arena was divided into 3 sections – one at each end representing TMA and BNZ, and a central “neutral” zone. The time spent in each zone was counted and a preference for either TMA or BNZ was determined.

### Stereotactic surgery

Stereotactic surgery was performed at least two weeks before starting any behavioral procedures. Mice were anaesthetized with isoflurane (3–5% for induction, 1–2% for maintenance) and head-fixed on the stereotaxic instrument. Ophthalmic ointment was applied to eyes in order to prevent drying and intracutaneous lidocaine was injected in the scalp. The surgical area was then washed with chlorhexidine and 70% ethanol three times before using a blade to perform a 1-1.5 cm incision through the midline of the scalp. After swabbing with increasing concentrations of ethanol, hydrogen peroxide was used to enhance visualization of lambda and bregma. Animals underwent bilateral craniotomy and holes were drilled at the appropriate coordinates for the target region, detailed below. The viral mix was injected using a micro syringe pump (UMP3; WPI) and a 10 µL Hamilton syringe (701LT; Hamilton) at the appropriate depth for the target region. Injection of 300 nL of virus was at a rate of 60 nL/min. After completion of the injection, the needle was left in the injection site for an additional 5 minutes. For optogenetic and calcium imaging experiments, the implant (Doric) was fixed to the stereotaxic adapter and was lowered in the brain, 0.1-0.2 mm above the virus injection site. To secure the implant, an even layer of Metabond was applied and let dry for about 10-15 minutes. Dental cement was then used to secure the cap to the implant. The protective cap was provided by cutting the top portion of a black polypropylene microcentrifuge tube. Animals were then left in a warm box (29°C) until recovered from anesthesia and then were returned to their homecage. On the day of the surgery and on the two following days mice were injected subcutaneously with meloxicam (10 mg/kg). Mice were allowed to recover for 14 days before any behavioral procedure was performed. Recovery was monitored with the use of a HPRA-approved recovery assessment table.

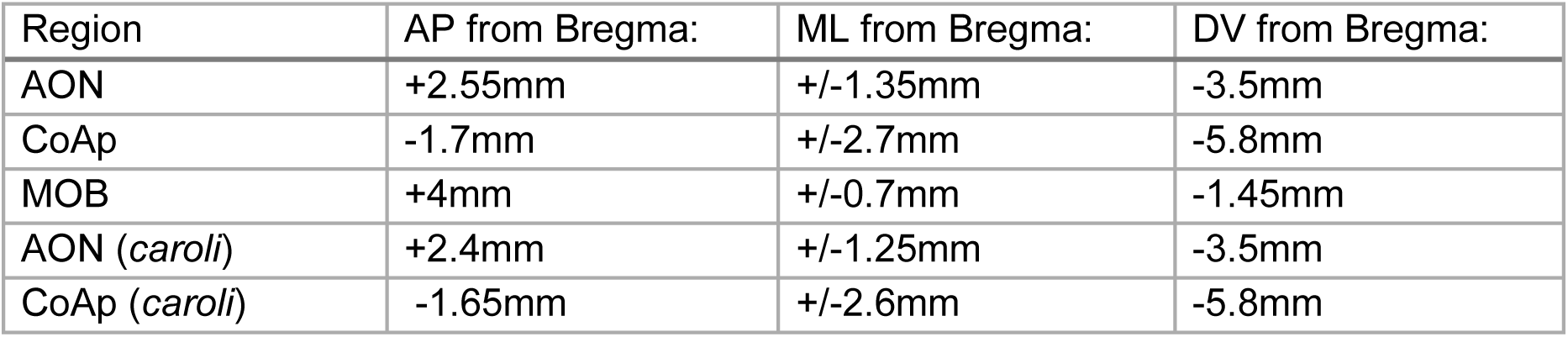

### Viruses

pAAV.Syn.GCaMP6f.WPRE.SV40 was a gift from Douglas Kim & GENIE Project (Addgene viral prep # 100837-AAV9; http://n2t.net/addgene:100837; RRID: Addgene_100837)

pAAV-hSyn-DIO-HA-hM4D(Gi)-IRES-mCitrine was a gift from Bryan Roth (Addgene viral prep # 50455-AAV8; http://n2t.net/addgene:50455; RRID: Addgene_50455)

pAAV-EF1a-double floxed-hChR2(H134R)-EYFP-WPRE-HGHpA was a gift from Karl Deisseroth (Addgene viral prep # 20298-AAV9; http://n2t.net/addgene:20298; RRID: Addgene_20298) pAAV-CaMKIIa-hM4D(Gi)-mCherry was a gift from Bryan Roth (Addgene viral prep # 50477-AAV9; http://n2t.net/addgene:50477; RRID: Addgene_50477)

| Virus | Addgene ref | Serotype | Titer | Dilution |
|---|---|---|---|---|
| AAV9-Syn-GCamp6f | #100837 | AAV9 | ≥ 1×10<sup>13</sup> vg/mL | 1 in 2 |
| pAAV-hSyn-DIO-HA-hM4D(Gi)-IRES-mCitrine | #50455 | AAV8 | ≥ 1×10<sup>13</sup> vg/mL | undiluted |
| AAV-EF1a-double floxed-hChR2(H134R)-EYFP-WPRE-HGHpA | #20298 | AAV9 | ≥ 1×10<sup>13</sup> vg/mL | 1 in 4 |
| pAAV-CaMKIIa-hM4D(Gi)-mCherry | #50477 | AAV9 | ≥ 1×10<sup>13</sup> vg/mL | undiluted |

### TRAP2 engram labelling and stimulation

TRAP2 mice underwent stereotactic surgery to express a Cre-dependent channelrhodopsin-2 in the AON or the CoAp. 2 weeks after surgery, mice were exposed to either clean air or TMA-odorized air for 5 min, followed by immediate intraperitoneal injection of 50 mg/kg 4-OHT. On the day of stimulation, mice were loaded into the chamber and the optical fiber implant was connected to a 450 nm laser diode fiber light source (Doric LDFLS 450/080). During the habituation session, the mouse was allowed to explore the chamber for 5 min, followed by a 5 min stimulation stage. During the light-on epochs, the mouse received blue light stimulation at 20 Hz with a pulse width of 10ms whenever the mouse crossed a boundary at a halfway point in the arena.

### TRAP2 engram inhibition

TRAP2 mice underwent stereotactic surgery to express a Cre-dependent inhibitory DREADD in the AON or CoAp. 2 weeks after surgery, mice were exposed to either clean air or TMA-odorized air for 5 min, followed by immediate intraperitoneal injection of 50 mg/kg 4-OHT. 2 weeks later, mice were injected intraperitoneally with either 2 mg/kg of clozapine-N-oxide in 0.9% w/v saline solution, or a saline control, 30 min prior to testing. Mice were then placed in the olfactory arena to assess their behavioral response to TMA. Mice were tested again 1 week later with the opposite treatment (saline or CNO).

### CaMKII inhibition in *Mus caroli*

CAROLI/EiJ mice underwent stereotactic surgery to express inhibitory DREADD in the AON or CoAp under control of the CaMKII promoter. 3 weeks after surgery, mice were injected intraperitoneally with either 2 mg/kg of clozapine-N-oxide in 0.9% w/v saline solution, or a saline control, 30 min prior to testing. Mice were then placed in the olfactory arena to assess their behavioral response to TMA. Mice were tested again 1 week later with the opposite treatment (saline or CNO).

### Immunohistochemistry

Mice were injected intraperitoneally with a terminal dose of Sodium Pentobarbital (50 µL) and perfused transcardially with PBS followed by 4% paraformaldehyde. Brains were then collected and left in 4% PFA at room temperature for 24 h, after which they were moved to PBS and stored at 4°C. Brains were cut using a vibratome into 50 µm coronal slices.

For the immunostaining, slices were first washed in PBT 0.2% three times for 10 minutes. Then, they were incubated in a blocking solution (PBT 0.2% + NGS 10%) for 1 hour at room temperature. They were incubated with primary antibody (in PBT 0.2% +NGS 3%) overnight at 4°C while shaking. On the following day, slices underwent three 10 min washes in PBT-0.2% before incubation with secondary antibody (PBT 0.2% + NGS 3%) for two hours in the dark at room temperature. Then, after three additional 10 min washes in PBT 0.2%, slices were stained with DAPI (1:1000 in PBS) for 15 minutes at room temperature. Slices were finally mounted on superfrost slides using Vectashield-DAPI. The antibodies used for the c-Fos staining were rabbit anti-c-Fos (Synaptic Systems #226008 at 1:1000) and anti-rabbit Alexa-568 (Invitrogen) (1:500). ChR2-EYFP was stained using a primary chicken anti-GFP IgY fraction (Invitrogen) (1:500) and rabbit anti-chicken Alexa-488 IgG (Invitrogen) (1:500).

### Image Acquisition and Analysis

Images were taken on a laser confocal microscope (Leica SP8 gated STED) with a 20x magnification unless otherwise stated. At least 4 images were taken for each region of interest per mouse. When the implant was not targeted correctly into the region of interest the animal was excluded from subsequent analysis. For each region of interest, density of cells was estimated by sampling cell density in the region of interest to calculate the number of DAPI cells per µm^2^. The total number of cells in a region was calculated by multiplying this value by the total area of the region counted. The number of c-Fos positive cells was manually counted using ImageJ.

### Whole-brain light sheet imaging Tissue processing

Following a 5 min exposure to trimethylamine (TMA)-odorized or clean air, mice were returned to their home cage for 60 min and then perfused transcardially with PBS followed by 4% paraformaldehyde (PFA) in PBS. Brains were post-fixed in 4% PFA for 24 h at room temperature, transferred to PBS, and stored at 4 °C until processing. Intact whole brains were then cleared and immunolabelled for c-Fos (monoclonal anti-c-Fos Rt, Sysy 226 017, 1:1000 dilution) detected with anti-Rt AlexaFluor647 (1:500 dilution, ThermoFisher A-21247) and Arc (polyclonal anti-Arc Rb, Sysy 156 003, 1:1000 dilution) detected with anti-Rb AlexaFluor488 (1:500 dilution, ThermoFisher A-32731) following the iDISCO+ protocol (*16*).

### Imaging

Cleared brains were imaged on a Luxendo MuVi-SPIM light-sheet microscope. The sample was illuminated simultaneously from both sides through opposing Nikon CFI Plan Apochromat Lambda D 4× objectives (NA 0.20, 20 mm working distance), producing a single fused light sheet, and emitted fluorescence was collected through one Olympus XLFluor 4×/340 detection objective (NA 0.28), giving a voxel size of 1.46 × 1.46 × 3 µm. Arc immunolabelling was visualized with 488 nm excitation and c-Fos with 647 nm excitation. Whole-brain imaging was performed across eight groups (n = 6-9 per group): male *Mus musculus* on standard chow, male *M. musculus* on a doxycycline diet, female *M. musculus* on standard chow, and male *M. caroli* on standard chow, each exposed to either TMA-odorized or clean (H₂O vehicle) air.

### Image Processing and Analysis

Following acquisition, overlapping image tiles were stitched into a single whole-brain volume in BigStitcher (*41*) running under Fiji/ImageJ v1.54p (*42*). An initial coarse alignment was computed on data downsampled by a factor of 4 × 4 × 2 in x, y and z, and tile positions were then refined by interest-point-based registration. Registered tiles were fused using linear interpolation with average blending across tile overlaps to produce the final volume. Raw light-sheet volumes were registered to the Allen Mouse Brain Atlas ([CCFv3]) as previously described (*17*). Labelled cells were detected and assigned to Allen Atlas-defined regions using ClearMap 2.0 (*18*). Two human annotators labeled individual volumes, and ClearMap parameters were chosen using a parameter sweep to identify the best match between automated and manual counts. Regional cell density was expressed as the natural log of the number of detected cells per voxel, normalized to the total number of cells detected in that brain. Fold change between conditions was computed per region as the log₂ ratio of mean cell counts in TMA- to H₂O-exposed animals and compared using two-sided Welch’s t-tests. P values were adjusted across regions within each comparison using the Benjamini–Hochberg false discovery rate procedure.

### Urinary trimethylamine quantification by ¹H-NMR spectroscopy

Urine samples (20 µl) were mixed with phosphate buffer (100 mM) containing 10% D2O and 0.5 mM TMSP, and pH adjusted to pH 7.0. Samples were then centrifuged at 13,000 g for 5 minutes. One-dimensional ¹H spectra were acquired using a standard spin-echo pulse sequence with water suppression and excitation sculpting on a Bruker DRX 500 MHz NMR spectrometer. Data were calibrated with respect to the TMSP signal. Peaks corresponding to trimethylamine (TMA), trimethylamine-N-oxide (TMAO), taurine and creatinine were identified based on comparison with previously published chemical shift values (*43*). Urinary TMA levels were normalized to the creatinine peak within the same spectrum to control for variation in urine concentration (*44*).

### Calcium imaging behavioral procedure

Calcium imaging was performed with a 0.5 mm GRIN lens (Doric). During calcium imaging sessions, mice were placed in the olfactory arena and recorded across three phases: a 5 min habituation period with H2O only, a 30 min odor test in which TMA-odorized air was presented against benzaldehyde (BNZ)-odorized air, and a 30 min social test in which a social stimulus was presented against a novel object, with a different object used for each social stimulus.

### Calcium imaging analysis

For each cell, Suite2p output (F, Fneu) was used to compute a neuropil-corrected fluorescence ratio across the mouse’s entire recording: (F − Fneu)/Fneu. This was z-scored using its mean and standard deviation across the whole recording. To test whether stimulus identity could be decoded from AON population activity, calcium traces were divided into 5s chunks for each stimulus. A random forest classifier (scikit-learn, balanced class weights) was trained for each mouse to predict stimulus identity from these feature vectors. Chunks from the same bout were always kept together on the same side of the train/test split, since chunks from one bout are not independent of each other and mixing them across the split would artificially inflate accuracy. Class sizes were balanced by downsampling, and this train/test procedure was repeated 100 times per mouse with independent random splits; classifier performance was summarized as the average per-class recall across repetitions. An identical procedure using shuffled stimulus labels was run in parallel to generate a chance-level control for comparison.

## QUANTIFICATION AND STATISTICAL ANALYSIS

Statistical analysis was performed using the pingouin and scipy.stats packages on Python. Data was tested for normality using the Shapiro-Wilk test. An unpaired Student’s T-test was used to compare means of two independent groups that assume equal variance between groups, while the paired test was used for dependent groups. For data with more than two groups, an ANOVA was performed to determine if there was a significant difference. If so, and the data was normally distributed, post-hoc analyses were performed for pairwise comparisons. For data with multiple comparisons from the same subjects, False discovery rate correction via the Benjamini-Hochberg method was performed to calculate adjusted q values. Analyses were performed in Python 3 using pandas (*45*), NumPy (*46*), SciPy (*47*) and statsmodels (*48*), with figures generated in matplotlib and seaborn.

## Declarations and Acknowledgements

## Acknowledgements

We thank Tamara Boto, Mani Ramaswami, and past and present members of the Ryan and Dennis Labs for support and discussion. We thank Shane O’Mara and the Comparative Medicine Unit of Trinity College Dublin for enabling *Mus caroli* experiments. We thank the Janelia Anatomy and Histology support team, particularly Monique Copeland, Morgan Clarke, and Benjamin Foster for their support.

## Author Contributions

P.B.C. and T.J.R. conceived of the study and designed experiments.

P.B.C., Z.W., N.S., E.U. and Í.M.C. performed experiments and analyzed the data.

P.B.C. prepared all figures. P.B.C, E.D., and T.J.R. interpreted the findings. P.B.C. and T.J.R. wrote the manuscript.

P.B.C. and T.J.R. acquired funding support. T.J.R. supervised the project.

## Inclusion and Diversity

We support inclusive, diverse, and equitable conduct of research.

## Supporting information

**Fig S1.**
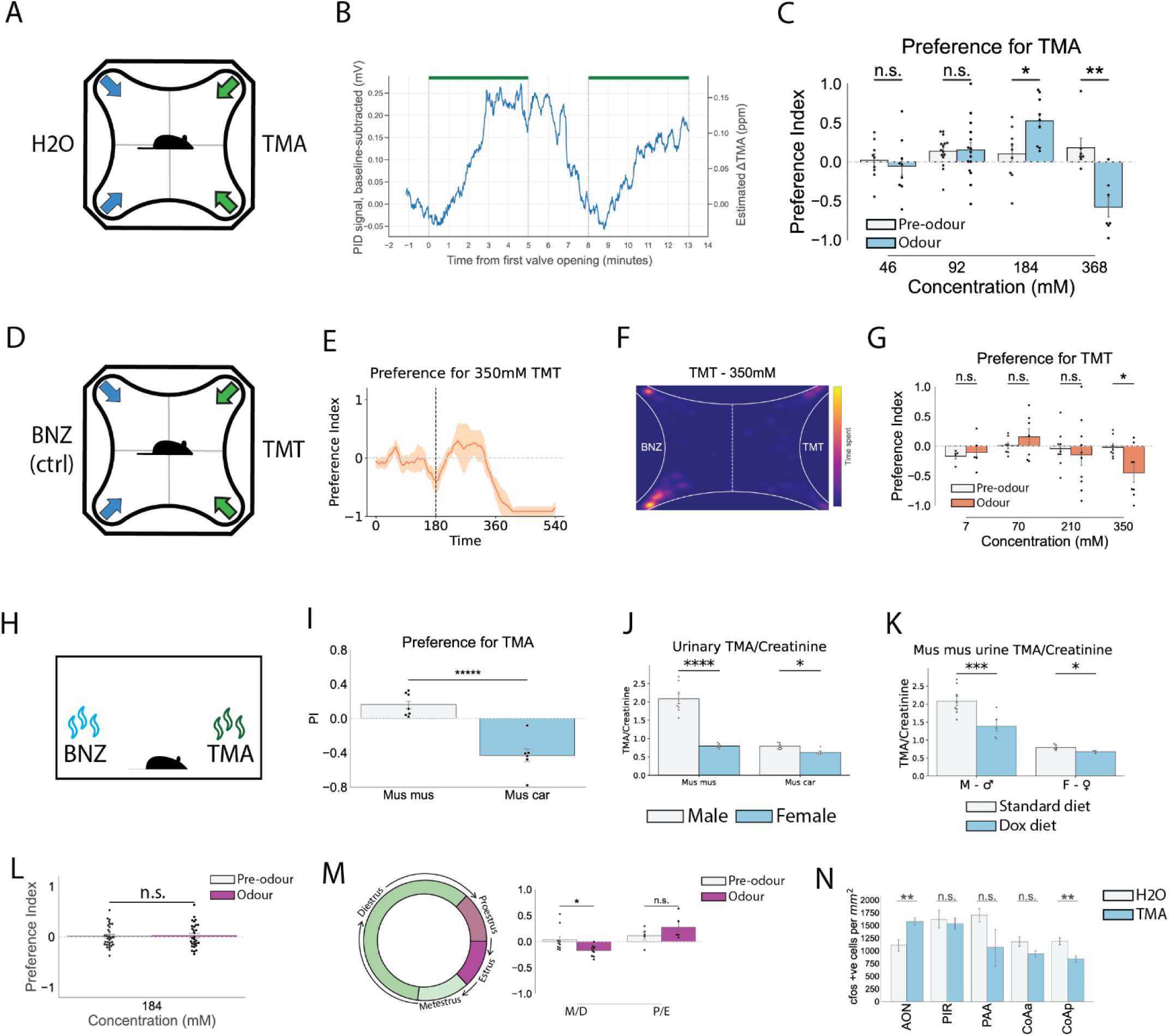
A vacuum-powered odor delivery system. Related to figure 1. (A) Schematic of vacuum-powered odor delivery apparatus. (B) Photoionization Device (PID) recordings indicating relative odor levels, with valve opening positions indicated by the green bars. (C) Response of *Mus musculus* to TMA over various concentrations (Student’s paired T test; 46 mM, n = 9; 92 mM, n = 15; 184 mM, n = 9; 368 mM, n = 7). (D) Schematic of vacuum-powered odor delivery apparatus for delivery of TMT. (E) Timetrace indicating how odor preference index evolves over time for 350 mM TMT (n = 8, shaded area represents standard error of the mean). Odor presentation indicated by a vertical dashed line. (F) Heatmap of TMT response. (G) Response of male *Mus musculus* to varying concentrations of TMT (Student’s paired T test; 7 mM, n = 5; 70 mM, n = 7; 210 mM, n = 10; 350 mM, n = 8). (H) Simple odor delivery apparatus, with 20 µL of odorant placed at each end of the rectangular arena. (I) Preference index of *Mus musculus* (n = 7) and *Mus caroli* (n = 6) to 184 mM TMA in the simple arena. (J) Urinary TMA levels in *Mus musculus* (male n = 8, female n = 5) and *Mus caroli* (male n = 6, female n = 5), normalized to creatinine. (K) Urinary TMA levels of Mus musculus on standard diet (male n = 8, female n = 5) and doxycycline diet (male n = 5, female n = 3). (L) Preference index of female *Mus musculus* mice towards 184 mM TMA (n = 30). (M) Preference index of female *Mus musculus* mice, separated by stage of the estrus cycle, towards 184 mM TMA (M/D n = 10, P/E n = 5). (N) Number of c-Fos positive cells detected on brain slices collected from male *Mus musculus* following exposure to H2O (n = 3) or TMA (n = 3). n.s. *p* ≥ 0.05, \**p* < 0.05, \*\**p* < 0.01, \*\*\**p* < 0.001, \*\*\*\**p* < 0.0001, \*\*\*\*\**p* < 0.00001

**Fig S2.**
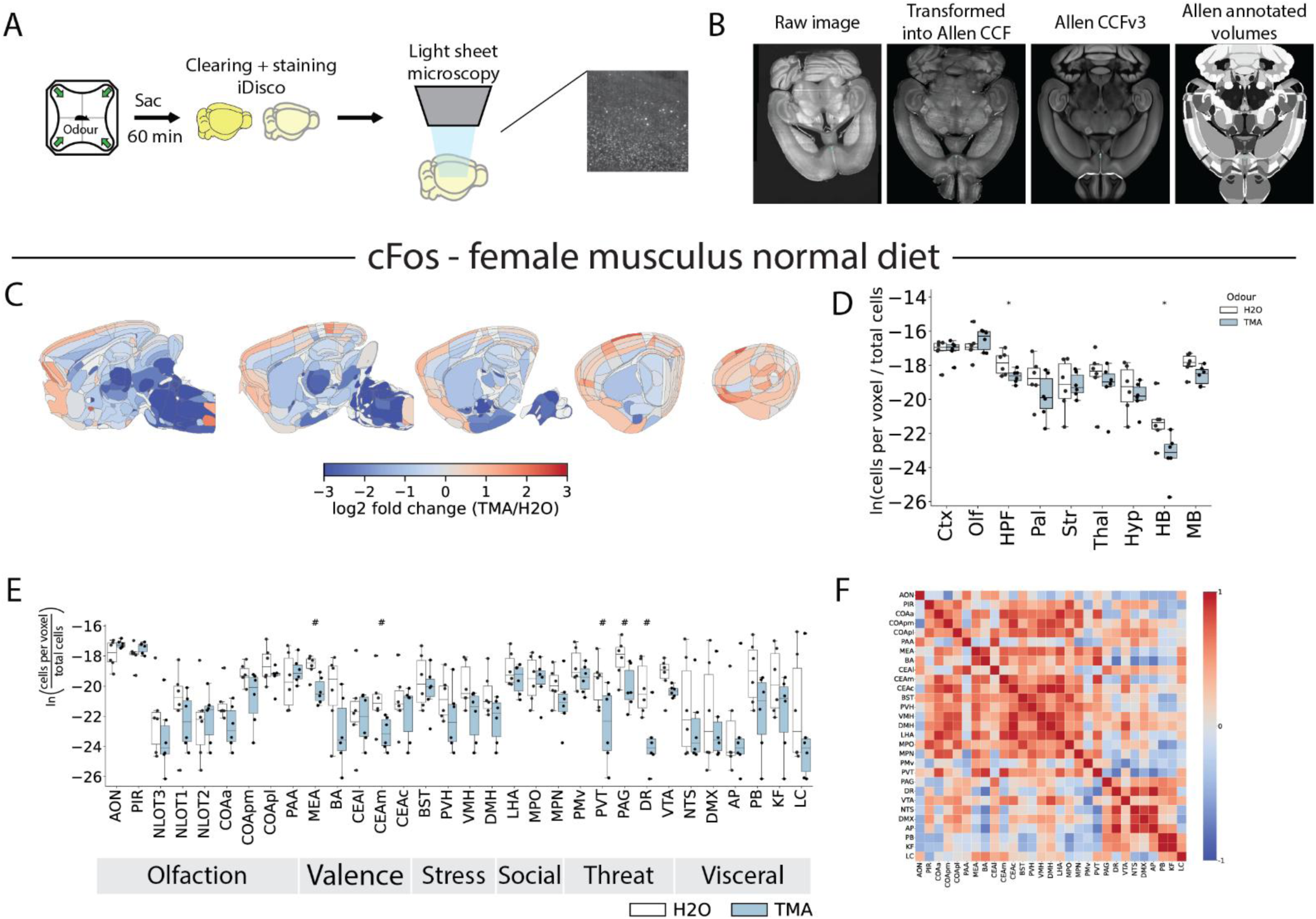
Reduced c-Fos expression in the hypothalamus and hindbrain in normal diet female *Mus musculus.* (A) Brains were cleared and stained according to the iDisco protocol and imaged on a light sheet microscope. (B) Raw images were transformed into a common coordinate framework and then fixed to the Allen Atlas template. Cells within each Allen Atlas region are quantified. (C) Fold change in c-Fos expression in the whole brain during TMA exposure (n = 6) vs. H2O exposure (n = 6) for female *Mus musculus* on a standard diet. (D) Cell counts for TMA and H2O exposed mice. (E) Relative c-Fos expression levels in selected brain regions during TMA vs. H2O exposure for female musculus mice on a normal diet. (F) Correlations of c-Fos expression across selected brain regions for TMA-exposed female *Mus musculus* mice on a normal diet. # *p* < 0.05, * q < 0.05

**Fig S3.**
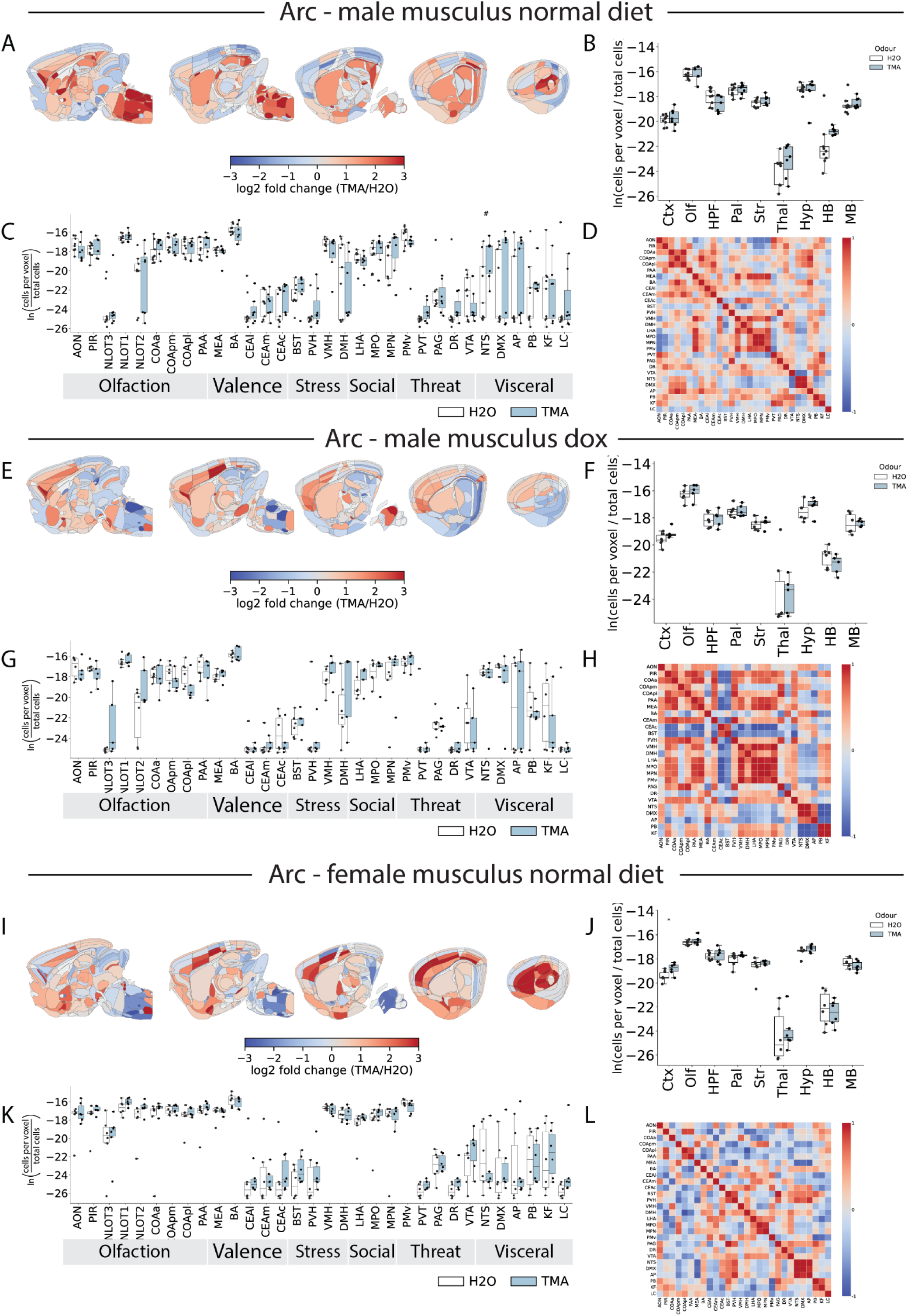
No large-scale changes to Arc expression in the whole brain of normal diet and doxycycline diet *Mus musculus.* (A) Fold change in Arc expression in the whole brain during TMA exposure (n = 7) vs. H2O exposure (n = 9) for male *Mus musculus* on a standard diet. (B) Cell counts for TMA and H2O exposed mice. (C) Relative Arc expression levels in selected brain regions during TMA vs. H2O exposure for male *musculus* mice on a normal diet. (D) Correlations of Arc expression across selected brain regions for TMA-exposed male *musculus* mice on a normal diet. (E-H) As in A-D, but for male *Mus musculus* on a doxycycline diet (H2O n = 6, TMA n = 6). (I-L) As in A-D, but for female *Mus musculus* on a normal diet (H2O n = 6, TMA n = 6). # *p* < 0.05, * q < 0.05

**Fig S4.**
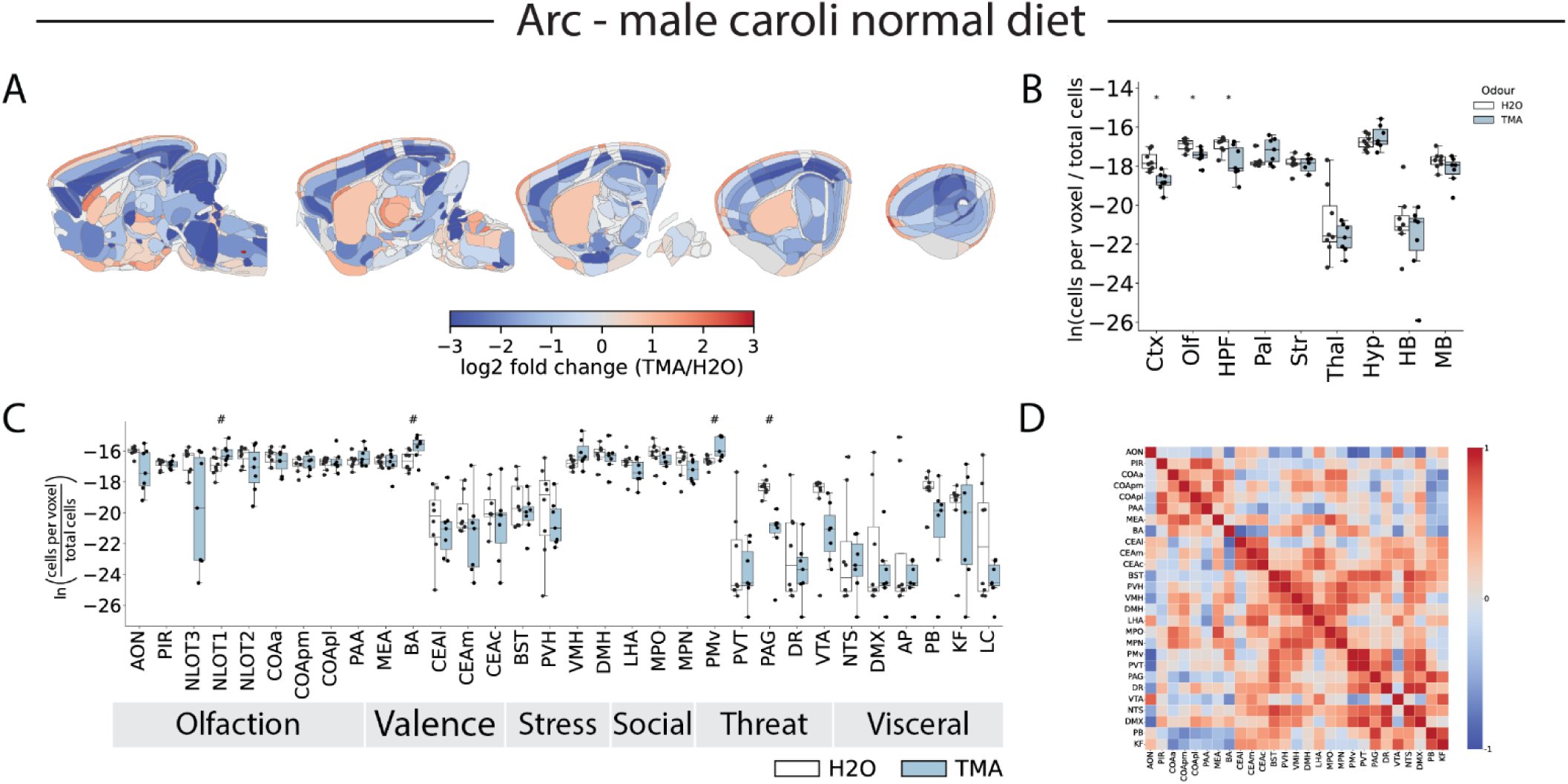
Decrease in Arc expression in the cortex in *Mus caroli.* (A) Fold change in Arc expression in the whole brain during TMA exposure (n = 7) vs. H2O exposure (n = 8) for male *Mus caroli*. (B) Cell counts for TMA and H2O exposed mice. (C) Relative Arc expression levels in selected brain regions during TMA vs. H2O exposure for male *Mus caroli* mice on a normal diet. (D) Correlations of Arc expression across selected brain regions for TMA-exposed male *Mus caroli* mice. # *p* < 0.05, * q < 0.05

**Fig S5.**
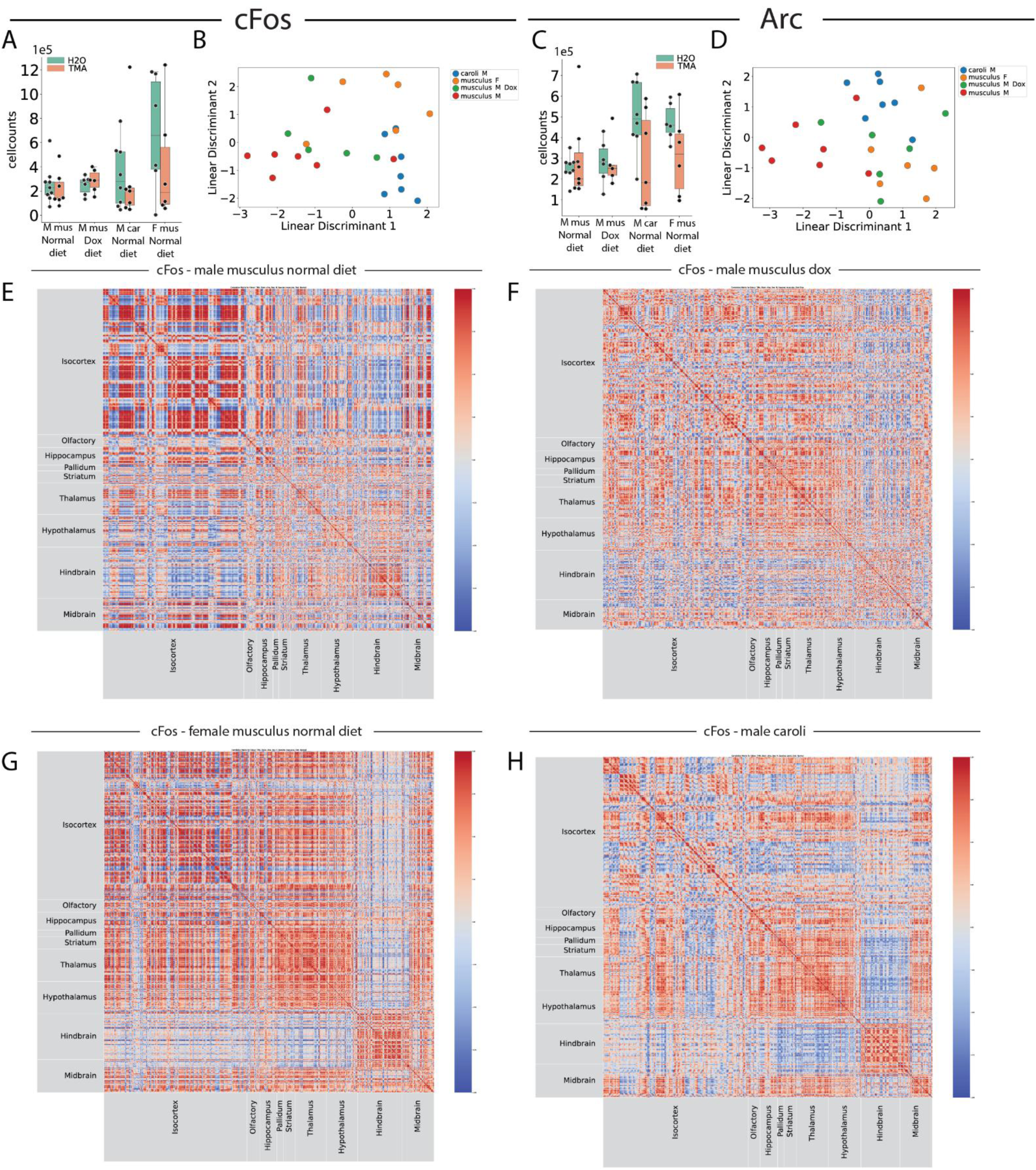
Correlations of whole brain c-Fos expression reveal distinct patterns within and between species. Related to figures 2 and 4. (A) Total c-Fos+ cells detected across each experimental group. (B) Linear discriminant analysis of whole brain c-Fos expression. (C-D) As in A-B, but for Arc+ cells. (E-H) Correlation matrices showing Pearson’s R correlation for the number of c-Fos+ cells detected across the whole brain for (E) male *Mus musculus* on a normal diet, (F) male *Mus musculus* on a doxycycline diet, (G) female *Mus musculus* on a normal diet and (H) male *Mus caroli* on a normal diet.

**Fig S6.**
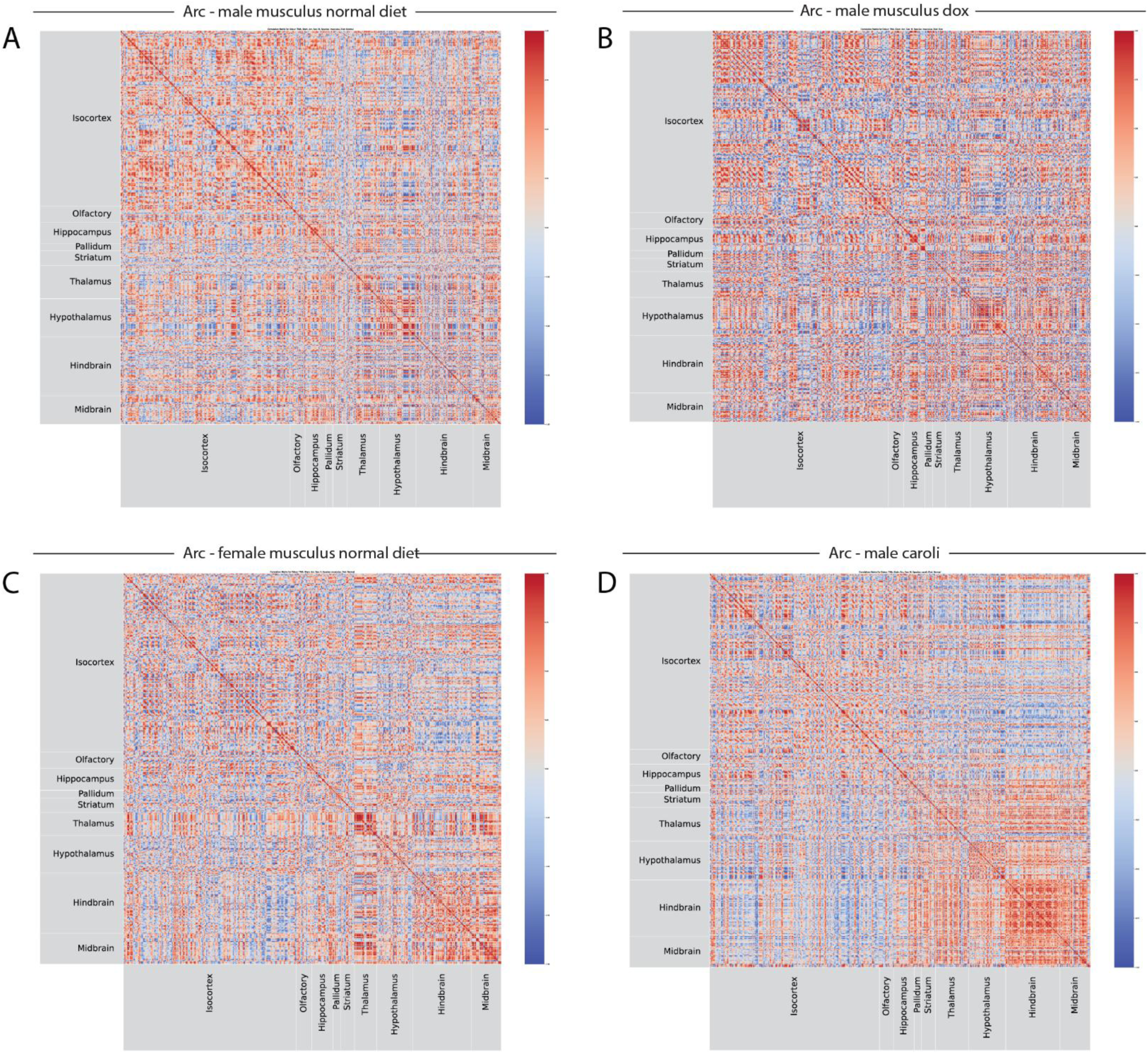
Correlations of whole brain Arc expression display subtle differences within and between species. Related to figures 2 and 4. Correlation matrices showing Pearson’s R correlation for the number of Arc+ cells detected across the whole brain for (A) male *Mus musculus* on a normal diet, (B) male *Mus musculus* on a doxycycline diet, (C) female *Mus musculus* on a normal diet and (D) male *Mus caroli* on a normal diet.

**Fig S7.**
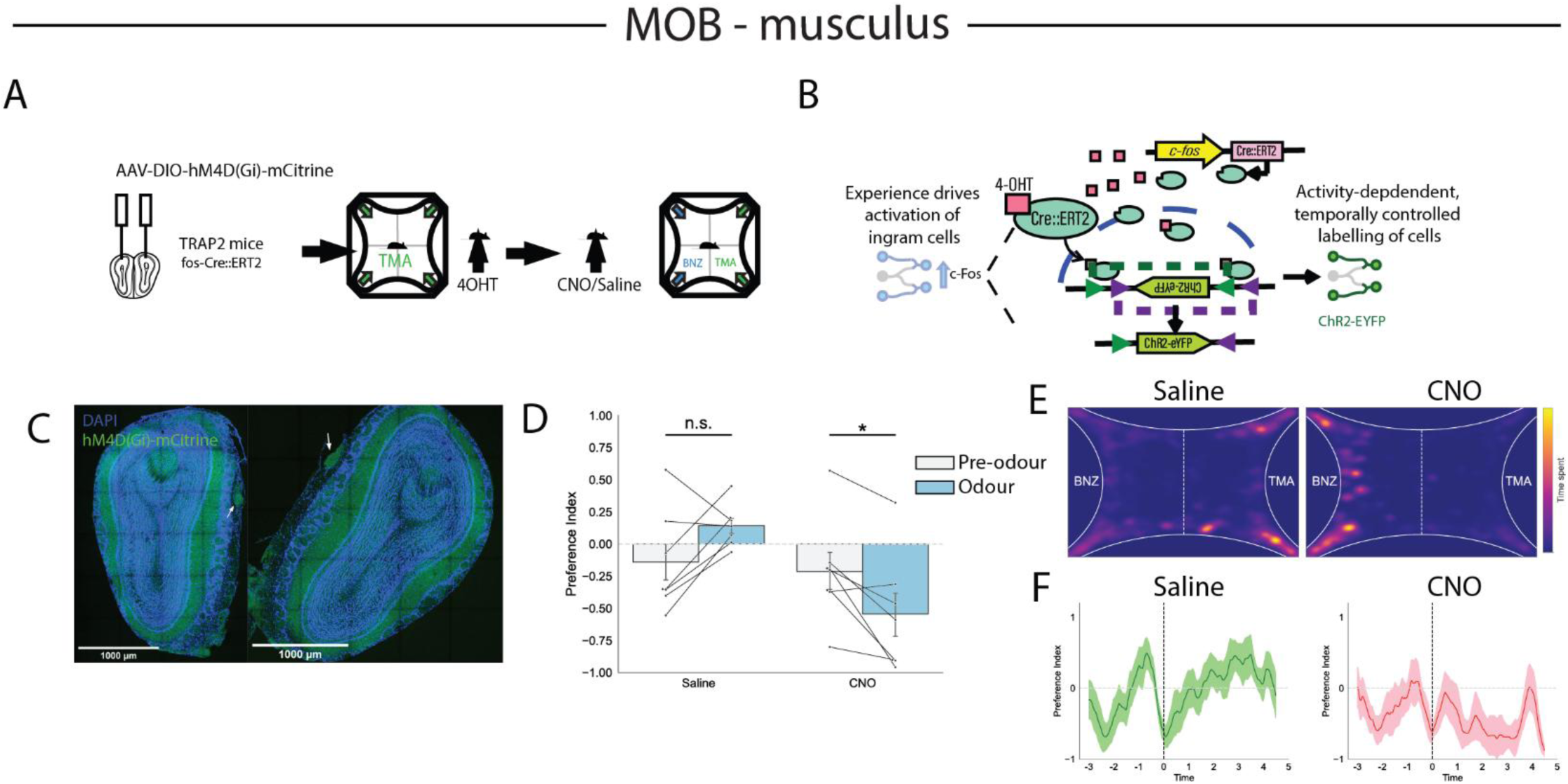
The main olfactory bulb is necessary for TMA attraction in *Mus musculus*. (A) Schematic indicating behavioral timeline for engram labelling and engram inhibition during TMA olfaction. (B) Activity-dependent cell labelling strategy. (C) Example slices showing expression of the inhibitory DREADD throughout the main olfactory bulb. (D) Preference index following administration of saline (n = 7) vs. CNO (n = 7). (E) Heatmaps of mouse position relative to odor location following saline or CNO administration. (F) Timetraces indicating average preference index across mice for saline or CNO administration. Odor onset indicated by dashed line. n.s. *p* ≥ 0.05, \**p* < 0.05

**Fig S8.**
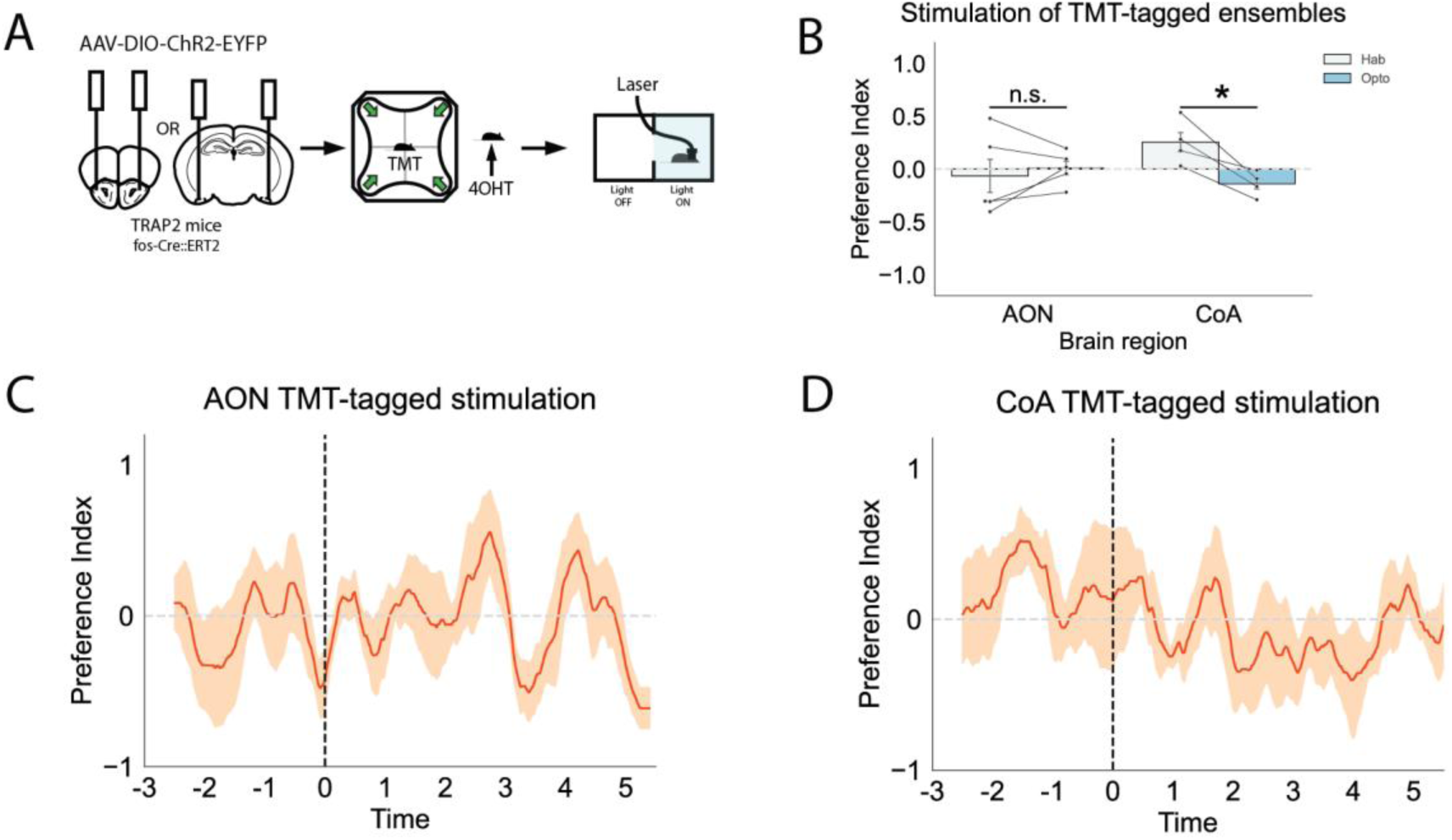
Stimulation of TMT-tagged cells in the CoAp but not the AON induces avoidance. (A) Schematic indicating behavioral timeline for engram labelling and spatially-controlled engram reactivation. Laser stimulation occurs in one half of the arena only. (B) Preference index pre- and post-onset of laser stimulation for AON (n = 5) or CoAp (n = 5) cells tagged during exposure to TMT. (C) Timetrace indicating average preference index across mice during optogenetic stimulation of AON cells tagged during delivery of TMT. Laser onset phase indicated by dashed line. (D) As in C, but for the CoAp. n.s. *p* ≥ 0.05, \**p* < 0.05

